# EMMAi: fast enzyme-allocation constraints in GEMs for improved biomass prediction across carbon sources

**DOI:** 10.1101/2025.05.01.651607

**Authors:** Juan P. Molina Ortiz, Derek Benson, James Watts, Mariana Velasque, Andrew C. Warden, Matthew J. Morgan

## Abstract

Genome-scale metabolic models (GEMs) predict emergent phenotypes by modeling the metabolic networks encoded in genomes. While GEMs have significantly advanced systems biology, metabolic engineering, biomedicine, and environmental science, they require extensive time and resources for manual curation, which can limit their utility in rapidly evolving research landscapes. Recent findings suggest that manually curated reactions can sometimes reduce prediction accuracy, indicating that integrating additional biologically grounded constraints may better capture emergent phenotypes. One promising approach is the incorporation of enzyme allocation constraints, which has been shown to enhance the predictive accuracy in metabolic models.

Enzymatically constrained GEMs (ecGEMs) rely on enzyme turnover rates (*kcat*) and protein molecular weights (MWs) to account for intracellular resource limitations by introducing an enzyme pool variable and assigning costs to reactions, thereby simulating enzymatic resource constraints. Tools such as GECKO, AutoPACMEN, and ECMpy provide computational pipelines for ecGEM generation. However, these pipelines are often limited by their reliance on experimentally measured kcat values or deep learning-predicted values, such as those generated by DLKcat, which face challenges in predicting kinetics for enzymes dissimilar to their training data. Additionally, these methods frequently require extensive manual curation of *kcat* values based on empirical data, a time-intensive process that hampers scalability and applicability to non-model organisms.

To address these limitations, we introduce EMMAi (Enzyme-constrained Metabolic Models with AI), a pipeline that fully automates the incorporation of enzyme constraints into GEMs. Unlike existing pipelines, EMMAi exclusively utilizes *kcat* values predicted by UniKP, an AI framework with improved accuracy over DLKcat, particularly for enzymes not present in training datasets. UniKP achieves a 13% improvement in correlation for unseen enzymes, enabling EMMAi to deliver ecGEMs with enhanced prediction accuracy without manual curation requirements.

We evaluated EMMAi by applying it to three GEMs: two manually curated models, *iJO1366* (Escherichia coli str. K-12 substr. MG1655) and *iMO1056* (Pseudomonas aeruginosa PAO1), and one draft GEM constructed and gap-filled using CarveMe. EMMAi-generated ecGEMs showed an average Pearson Correlation Coefficient (PCC) improvement of 0.27 for manually curated GEMs when compared to predicted and experimentally measured growth rates and Biolog readings. Notably, for the draft GEM of *Pseudomonas aeruginosa PAO1*, the PCC improved dramatically from −0.3 to 0.6.

EMMAi demonstrates that automating the integration of enzyme allocation constraints using AI-predicted kinetic parameters significantly enhances the prediction accuracy of GEMs, even in the absence of manual curation. These results underscore EMMAi’s potential as a scalable, efficient, and accurate tool for advancing GEM-based research in systems biology, metabolic engineering, and beyond.

## Introduction

Genome-scale metabolic models (GEMs) aim to predict emergent phenotypes by modelling the metabolic network encoded in genomes while simplifying or abstracting other aspects of the cellular system (1). Metabolic reactions in GEMs are represented as linear equations arranged in a stoichiometric matrix (*S*-matrix), allowing for the calculation of flux distributions. Through the application of constraint-based analyses (CBA), such as flux balance analysis (FBA) (2, 3), they have been widely used to inform the fields of systems biology (4, 5), metabolic engineering (6, 7), biomedicine (8, 9) and environmental science (10, 11).

With technology and biological research evolving at an unprecedented pace, there is a pressing need for the development of methodologies capable of producing outcomes that can match that tempo. Meanwhile, GEMs still demand important time and resource investment for evidence-based manual curation (12). Moreover, it has been recently reported that addition of manually curated reactions to GEMs of the model organism *E. coli*, can hinder prediction accuracy rather than improving it (13). This suggests that conceptualising additional biologically grounded constraints into GEMs might represent a better alternative to capture emergent phenotypes more accurately. Among these, efforts to incorporate enzyme allocation constraints to conventional GEMs have shown promise.

Enzymatically constrained GEMs (ecGEMs) have stricken the right balance between improving growth rate prediction accuracy and assuring that the modified GEMs remain compatible with most constraint-based methods and platforms (14–17). Computational pipelines such as GECKO (14), AutoPACMEN (15) and ECMpy (17) allow the incorporation of enzyme constraints to conventional GEMs from experimentally measured and/or predicted enzyme turnover rates (*kcat* values) and protein molecular weights (MWs). AutoPACMEN retrieves experimentally measured *kcat* values from the online databases BRENDA (18) and SABIO-RK (19), limiting its application to GEMs of model organisms or closely related strains. While GECKO (14) and EMCpy (17) can also source *kcat* values from these online databases, DLKcat (20), a deep learning algorithm for *kcat* prediction, has been adapted to their latest versions, theoretically expanding their applicability to non-model organisms. However, DLKcat has been under strong scrutiny due to its limitations to predict accurate *kcat* values for enzymes dissimilar to its training data (21). Furthermore, currently available algorithms either need or strongly recommend manual or semi-automated adjustment of *kcat* values based on empirical data (14, 15, 17), which can result in an extremely time-consuming exercise. Therefore, an improved ecGEMs implementation pipeline, that offers better phylogenetic coverage and minimises manual curation requirements, is needed.

To address this need, we developed EMMAi (Enzyme-constrained Metabolic Models with AI), a pipeline to automatically incorporate enzyme allocation constraints into conventional GEMs. EMMAi follows a similar methodology as GECKO and AutoPACMEN, where the S-matrix in conventional GEMs is modified to incorporate an *enzyme cost* pseudo-metabolite in each reaction, with a specific *cost* coefficient derived from enzyme molecular weight (MW) and corresponding kcat value, and an *enzyme pool* pseudo-reaction. These additional components constrain the total flux in a GEM by conceptualising limited availability of intracellular resources for the synthesis of enzymes. However, EMMAi relies exclusively on *kcat* values predicted from UniKP (22), an AI structure for the prediction of enzyme kinetics, which reports a 13% correlation improvement over DLKcat for enzymes not present in their training set. This characteristic enables EMMAi to produce ecGEMs with a significatively higher accuracy across carbon source growth predictions, without the need of significant postprocessing or constraint recalibration.

To demonstrate the improvements in prediction accuracy in ecGEMs built with EMMAi, we implement three enzymatically constrained models from conventional GEMs. Two manually curated GEMs (iJO1366, *Escherichia coli* str. K-12 substr. MG1655 (23) and iMO1056, *Pesudomonas aeruginosa* PAO1 (24)) and one automatically generated and gapfilled utilising CarveMe (25). Overall, we show that EMMAi improves Pearson Correlation Coefficient (PCC) between predicted and experimentally measured growth rates and Biolog readings by 0.27 on average for manually curated GEMs. Further, we find an improvement in PCC of approximately 0.9 (from −0.3 to 0.6) in a *Pseudomonas aeruginosa* PAO1 draft GEM built and gapfilled with CarveMe.

## Methods

### Computational resources

#### Software Framework

EMMAi was developed using Python 3.9 for compatibility with scikit-learn version dependencies. Dependencies are managed and distributed by the Conda package and environment management system. An installation of Conda or Mamba is recommended for environment management for local runs. EMMAi is also compatible with High Performance Computing (HPC) and job scheduling and management systems (e.g. SLURM). Access to a GPU cluster is recommended. As inputs, EMMAi requires a conventional GEM file in SBML or XML format, an optional protein FASTA file and a custom YAML file populated with additional GEM and genome data. EMMAi’s outputs include an enzymatically constrained GEM, as well as intermediate data files including protein sequence data and metabolite SMILES-sequence pairs data. Intermediate GEMs files are generated during this process.

### Model acquisition and construction

#### *Pseudomonas aeruginosa* PAO1 models

The manually curated GEM of *Pseudomonas aeruginosa* PAO1, iMO1056 was obtained from (24), which additionally reports on experimental growth in 31 carbon sources in defined medium.

For the assembly of a draft *Pseudomonas aeruginosa* PAO1 GEM we utilised the command line tool CarveMe (25). The protein FASTA file required for this procedure was downloaded from NCBI (26) for the corresponding organism under the identifier GCF_000006765.1. To reconcile growth/no growth outcomes across the 31 tested carbon sources, gapfilling was performed. We utilised the *– gapfill* function from CarveMe to address false negatives (no growth predicted by the model when experimental data from (24) reported growth). This process was limited to carbon sources with a metabolite identifier in BiGG (27), the reaction and metabolite database utilised by CarveMe. Files required to reproduce this GEM, including the specific commands submitted to CarveMe and the resulting SBML file, can be found in Supplementary file 3.

#### *Escherichia coli str*. K-12 *substr*. MG1655 models

The original *Escherichia coli str*. K-12 *substr*. MG1655 GEMs iML1515, and its predecessor iJO1366, were downloaded from the BiGG models database (27). The AutoPACMEN version of iJO1366 before calibration was obtained from the data repository in (15) – model name: iJO1366_sMOMENT_2019_06_25_STANDARD_EXCHANGE_SCENARIO.xml.

### Modelling media and growth predictions

Media defined in (15) for *Escherichia coli str*. K-12 *substr* growth and for modelling their ecGEM was used for models derived from iJO1366 with no carbon sources. Meanwhile, iMO1056 and the derived ecGEM were modelled in the media defined in (24) without glucose. The draft PAO1 GEM and ecGEM were modelled utilising a modified version of the media in (24) and the M9 media defined in CarveMe (25) without glucose.

Growth predictions for each ecGEM were generated with no explicit limits (bounds, uptake rates) for individual carbon sources. Such rates are instead dependent on enzyme constraints, as done previously (15). The protein pool size for the EMMAi iML1515 ecGEM was adjusted to match the measured growth rate in glucose, as reported in (28). The size of the protein pool for both *P. aeruginosa* ecGEMs was calibrated to match the predicted growth rate of iMO1056 in glucose (24).

Experimental growth rates for benchmarking modelling predictions for *E. coli* were obtained from (28) and Biolog measurements reported in (24) were utilised to assess prediction accuracy in *P. aeruginosa* models. GEMs and ecGEMs were modelled with FBA utilising the GEMs modelling platform COBRApy (29) and the Gurobi solver (30). The modified GEMs, their details and the corresponding scripts are provided in Supplementary files 1, 2 and 3. A link to the repository with the command-line version of EMMAi will be added to this preprint once it is released.

### Scatterplots, linear regression and correlation estimations

We assessed the fit between model-predicted and experimentally measured growth rates using linear regression analysis and Pearson correlation tests. Experimental data included (i) optical density (OD_600_) measurements, representing direct growth rates, and (ii) Biolog respiration activity readings, which reflect metabolic activity across different carbon sources. For each metabolic model, we calculated the Pearson correlation coefficient (r) to evaluate the strength and direction of the relationship between predicted and observed growth rates. Statistical significance was assessed using p-values. All analyses were conducted in R (version 4.4.2) using the ggplot2, dplyr, and tidyverse packages.

## Results

### Algorithm

EMMAi incorporates proteome enzyme allocation constraints to conventional GEMs in two stages. A data collection stage (Stage 1), where EMMAi collects the data required to enzymatically constrain a GEM, and a model modification stage (Stage 2), where the structure of the GEM is modified, and the data collected is utilised to enzymatically constrain reactions in the model.

#### Stage 1 – data collection

Three dimensions of data are required to produce an ecGEM. Enzyme constraints are derived from two variables – molecular weights (MWs) and *kcat* values (Figure 1). Meanwhile, *kcat* values are predicted from substrate SMILES and the protein sequence associated to the corresponding reaction. Therefore, during this stage, EMMAi extracts data from the base GEM (reaction, metabolite and gene data), PubChem and/or ChemSpider (metabolite SMILES), and either a protein fasta file or UniProt (protein sequences and MW). When a protein fasta file is provided EMMAi employs Biopython to calculate the MW of protein sequences. When a fasta file is not available, it is necessary for gene IDs in the original GEM to be gene or protein identifiers compatible with UniProt as EMMAi will retrieve the corresponding sequences from this database, based on matches at the strain or species level, which need to be specified by the user. For the collection of metabolite specific SMILES, EMMAi utilises metabolite names in the original GEM to query PubChem and ChemSpider. Once the data collection process has been completed, EMMAi feeds protein sequence-SMILES pairs to UniKP for the prediction of *kcat* values. Protein sequence-SMILES pairing is based on protein-reaction associations in the GEM as well as reaction stoichiometry and directionality. Reaction cofactors and common co-reactants (e.g. H2O) are not considered for this pairing process as EMMAi aims to only pair protein sequences with main reaction substrates. Importantly, EMMAi does not consider transporter sequence-SMILES pairs for this process, as it would be biologically inaccurate to constrain transporters with a *kcat* value. This is the most time-consuming step in the EMMAi pipeline for a CPU based system (59 minutes on an Intel(R) Xeon(R) Platinum 8358 CPU @ 2.60GHz, 16 Core(s)).

**Figure 1.**
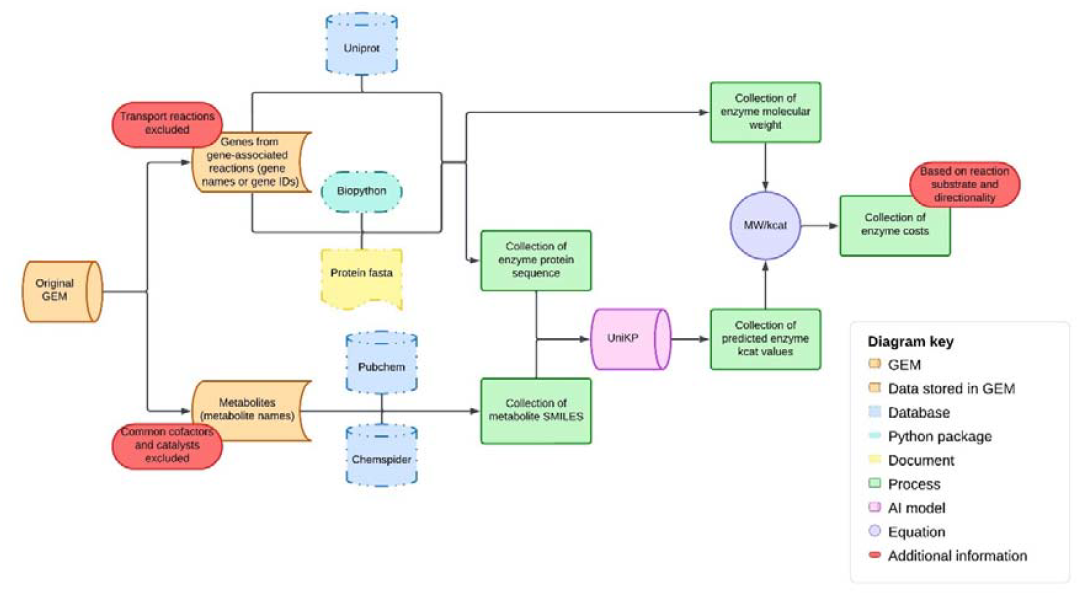
Data collection stage. Before implementing enzyme constraints into a conventional GEM, EMMAi automatically collects the data required to produce an ecGEM. This includes protein sequences, molecular weight (MW), metabolite SMILES from various sources and utilises UniKP to predict kcat values.

For a GPU-based system, the time is significantly improved (90sec when using 4x NVIDIA H100 GPUs).

#### Stage 2, model modification

Once the necessary *kcat* values have been predicted by UniKP, the original GEM can be modified to include the corresponding enzyme constraints or costs (Figure 2) which are calculated with the following equation:

**Figure 2.**
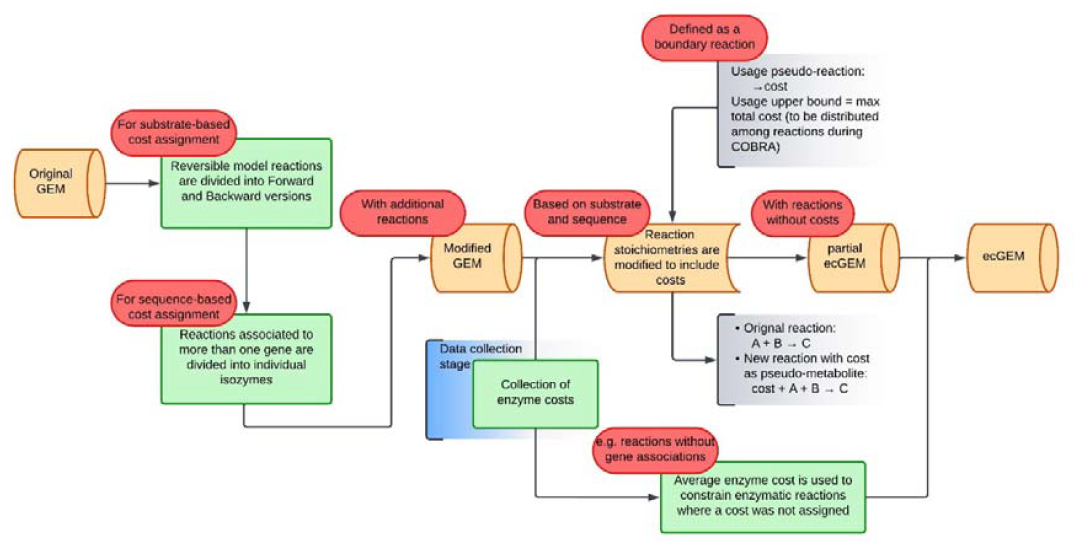
Model modification stage. Once the data required to produce an ecGEM has been collected and processed, EMMAi introduces a series of modifications to the base (conventional) GEM which modifies the S matrix by introducing an enzyme cost pseudo-metabolite and a protein pool pseudo-reaction, which limits the availability of the enzyme cost pseudo-metabolite.

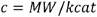

Where *c* is the cost coefficient required for a given reaction to run fluxes, calculated from relationship between the molecular weight (*MW)* of the enzyme catalysing the reaction and its *kcat* value. The rationale for the above equation, which is also applied by AutoPACMEN, GECKO and ECMpy, is centred around the assumption that cells select for the most resource-efficient pathways when expressing a given phenotype. In ecGEMs, this decision process is abstracted into two factors, MW and *kcat value*, arguing that enzymes with a low MW and a high *kcat* value are more resource-efficient options. To correctly estimate this relationship, reversible reactions, with different substrate, and therefore different *kcat* value, in each direction, must be divided into their corresponding forward and backward versions. This also means that reactions that are associated to more than one gene or gene complex (isozymes), must be divided into an equal number of individual reactions, as the corresponding MW and predicted *kcat* value will be different for each of them. In reactions with more than one main substrate, the lower predicted (rate-limiting) *kcat* value is selected. For reactions associated with a gene complex, the predicted *kcat* values are averaged.

To integrate the resulting costs into GEMs, while keeping their structure compatible with conventional COBRA platforms, modification of its *S* matrix is required. A cost pseudo-metabolite and a protein pool pseudo-reaction are incorporated into such matrix where the cost pseudo-metabolite is added to the righthand side of every enzymatic rection (substrates) where its coefficient is equal to the reaction’s calculated cost (MW/kcat). Enzymatic reactions for which a cost could not be determined (e.g. reactions without a gene association) are assigned a cost equals to the average of the calculated costs. Importantly, other non-biological reactions, such as biomass reactions, non-growth associated metabolism (NGAM), and boundary reactions, remain unmodified and costless (unconstrained). Meanwhile, the protein pool pseudo-reaction is added as a boundary reaction, in this case a demand reaction, for the cost pseudo-metabolite, where its lower bound represents the theoretical maximum amount of enzymogenic resources in the cell. This is given as the fractional weight of such resources per gram of dry weight (g/gDW).

Once the model modification process is completed the size of the protein pool pseudo-reaction (lower bound) needs to be adjusted so that the predicted growth rate in a given media formulation approximates its empirically measured counterpart. Once this process is completed, flux distributions predicted through CBA are going to be limited by the size of the protein pool pseudo-reaction, which has to be distributed across the represented enzymatic reactions that carry flux proportionally (e.g. higher fluxes will demand a higher proportion of the available cost pseudo-metabolite from the protein pool).

### Comparing EMMAi to preexisting pipelines

While EMMAi shares notable similarities with previously available ecGEM-building pipelines, key characteristics make of it a more suitable alternative (Table 1, Figure 3). EMMAi follows a similar approach to GECKO 3.0 and AutoPACMEN, where the *S*-matrix in a given GEM is modified to incorporate enzyme costs as pseudo-metabolite coefficients and a protein pool pseudo-reaction, that limits the availability of the enzyme cost pseudo-metabolite. However, EMMAi presents key methodological advantages that lead to improved cost calculation and therefore, improved phenotype predictions, in a reduced period of time. EMMAi utilises UniKP for *kcat* value predictions, which has shown improved performance over DLKcat (21), used by GECKO and ECMpy. Additionally, alternative pipelines can retrieve *kcat* values from the online databases BRENDA and/or SABIO-RK. AutoPACMEN relies on data stored in these resources exclusively, which is mostly limited to kinetics of model organisms. Yet, such functionality is not required when running EMMAi, given that UniKP can predict such values with high accuracy, as it was trained with enzyme sequences from both databases (22).

**Table 1.** Comparison of the main characteristics of EMMAi against pre-existing ecGEMs-building pipelines.

| **Characteristics** | **AutoPACMEN / sMOMENT** | **GECKO 3.0** | **ECMpy 2.0** | **EMMAi** |
| --- | --- | --- | --- | --- |
| Coding language | Python and MATLAB (model calibrator) | MATLAB | Python | Python |
| Method for resource allocation | Pseudo-metabolites (enzyme costs) and a protein pool pseudo-reaction integrated into the S-matrix | Pseudo-metabolites (enzyme costs) and a protein pool pseudo-reaction integrated into the S-matrix | A total protein pool constraint not integrated into the S-matrix | pseudo-metabolites (enzyme costs) and protein pool |
| Automation | Semi-automated (sequential scripts) | Minimal automation | Semi-automated (sequential Jupyter notebooks) | Fully automated (command-line) |
| *Kcat* value prediction model | Function not available | DLKcat | DLKcat | UniKP |
| Access to public databases for *Kcat* value retrieval | BRENDA, SABIO-RK, custom DB based on substrate name and EC number | BRENDA, custom DB based on substrate name and EC number | BRENDA, SABIO-RK, custom DB based on substrate name and EC number | Not required – estimated de novo |
| Protein sequences and molecular | UniProt | UniProt | UniProt | UniProt, FASTA files and |

| **Characteristics** | **AutoPACMEN /**<br>**sMOMENT** | **GECKO 3.0** | **ECMpy 2.0** | **EMMAi** |
| --- | --- | --- | --- | --- |
| weight |  |  |  | Biopython |
| SMILES | NA | PubChem | PubChem | PubChem,<br>ChemSpider,<br>VMH metabolite<br>database |
| Parameter<br>calibration | Script aided<br>manual curation<br>and<br>AutoPACMEN<br>model calibrator:<br>maximal kcat<br>value change<br>factor based on<br>kcat value<br>deletion impact | Semi-automated<br>iterative<br>refinement:<br>identification of<br>enzymes<br>overconstraining<br>the model<br>(demanding the<br>largest fraction of<br>the protein pool) | Iterative<br>refinement:<br>relaxation of Kcat<br>values in a given<br>pathway based on<br>max Kcat value<br>associated to the<br>same EC number<br>in BRENDA or<br>SABIO-RK | Not required |
| Parameter<br>optimisation time<br>(reported) | 24 hours +<br>manual curation<br>(time not<br>specified) | 15 min +<br>comparison<br>against literature<br>recommended<br>(time not<br>specified) | Not specified | Not required |

**Figure 3.**
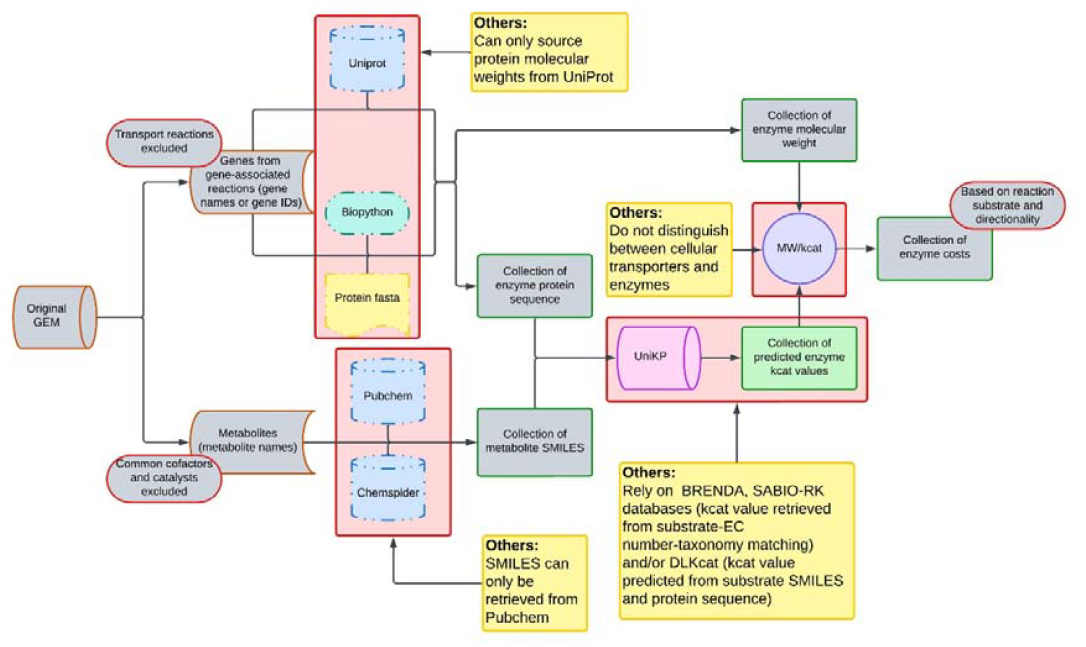
Differences between EMMAi and currently available pipelines for the implementation of ecGEMs. The main differences between EMMAi and alternative ecGEMs building pipelines is highlighted in a red background, together with a description in yellow.

*Kcat* value predictions from UniKP and DLKcat require substrate SMILES and protein sequences. While GECKO and ECMpy allow SMILES retrieval from PubChem, EMMAi expands this process to include ChemSpider, offering improved coverage of the metabolites in a given metabolic network. The four pipelines we compare in Table 1 can retrieve protein sequences and protein MW from UniProt through protein identifiers compatible with the platform. However, increased access to sequencing technologies and pipelines for the rapid assembly of GEMs has enable the construction of metabolic models of non-model organisms, for which protein identifiers are often not available in UniProt. Therefore, EMMAi can read protein fasta files for protein sequence retrieval, where standardised identifiers are not required. In such cases, MWs for individual sequences are calculated utilising the Python package Biopython.

Due to a limited access to reliable *kcat* values, the pipelines we compare EMMAi against, have developed custom approaches for calibrating individual enzyme costs, mostly due to restrictive *kcat* values (14, 15, 17). However, due to the application of UniKP and the unique features in EMMAi, the time expensive process of selective *kcat* value relaxation is not required for the generation of ecGEMs with improved prediction accuracy across dozens of carbon source, as we show in the following section.

### EMMAi significantly improves experimental-to-predicted growth rate correlations

To demonstrate the practical improvements our pipeline offers over preexisting approaches, we implemented enzyme constraints with EMMAi to manually curated models of *Pseudomonas aeruginosa* PAO1 and *Escherichia coli str*. K-12 *substr*. MG1655, as well as to an automatically built draft GEM of PAO1. We find that the enzyme constraints implemented by EMMAi lead to significant improvements in prediction accuracy, when correlated with experimental measurements, even before the application of *kcat* value recalibrations.

#### Pseudomonas aeruginosa PAO1

A manually curated P. aeruginosa PAO1 model, iMO1056 (24), was used as reference to build and ecGEM with EMMAi (Methods). Each model was simulated across 31 carbon sources, and the resulting growth predictions were correlated against previously reported Biolog growth estimations (24). We find that Pearson Correlation Coefficient (*r*), improves by 0.31 (from 0.36 to 0.67) in the EMMAI ecGEM (Figure 4A).

**Figure 4.**
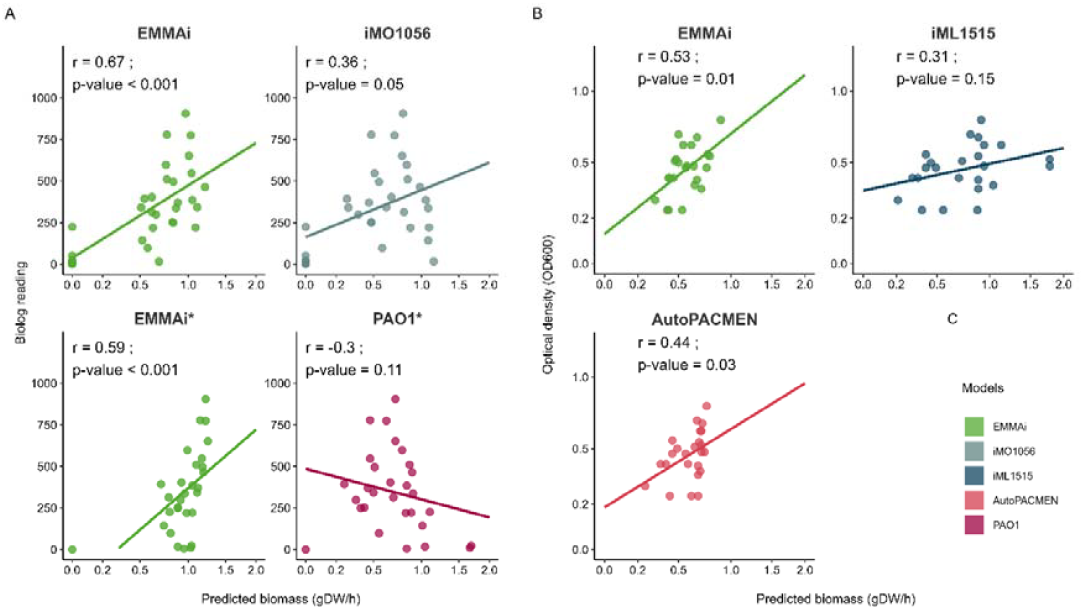
Incorporating enzyme constraints to manually curated and automatically generated GEMs improves growth rate prediction correlation across carbon sources. A) Comparison between growth rate predictions in a manually curated GEM (iMO1056, upper left, EMMAi constrained iMO1056, upper right) and a CarveMe generated GEM (base GEM, lower left, EMMAi constrained GEM, lower right) of Pseudomonas aeruginosa PAO1 and their correlation (r) with experimental (Biolog readings) results across 31 carbon sources. B) Comparison between growth rate predictions in a manually curated GEM (iJO1366, upper left), an ecGEM built with AutoPACMEN (iML1515, upper right) and a GEM constrained with EMMAi (lower left) of Escherichia coli str. K-12 substr. MG1655 and their correlation (r) with experimental results (OD) across 24 carbon sources. * Automatically generated Pseudomonas aeruginosa PAO1 GEM.

An increased number of research efforts have recently opted to obtain metabolic insights from automatically built and minimally curated GEMs, utilising publicly available resources such as CarveMe (25). Therefore, we applied EMMAi to a conventional PAO1 GEM automatically built with CarveMe and compared predictions derived from both models against Biolog growth estimations. Our analysis reports a 0.9 *r* improvement, from −0.3 in the draft GEM to 0.6 in the EMMAi ecGEM (Figure 4A).

#### Escherichia coli str. K-12 substr. MG1655

An ecGEM of the latest iteration of the Escherichia coli str. K-12 substr. MG1655 GEM, iML1515 (27), was created utilising EMMAi and the predictions from both were correlated with experimental growth measurements across 24 carbon sources (28). Such comparison reveals that the EMMAi implemented ecGEM improves such correlation by 0.22, when compared to the base model iML1515 (Figure 4B). We additionally compared these predictions with the version of the ecGEM built utilising AutoPACMEN, before semi-automated kcat value calibration. The EMMAi produced ecGEM was found to produce predictions that better correlate with experimental measurements (0.53 vs 0.41).

## Discussion

Here, we introduce EMMAi, a pipeline to enzymatically constrain conventional GEMs and improve growth rate predictions with no manual curation. Enzymatic constraints lead to more biologically accurate GEMs by incorporating finite protein resources and enzyme kinetics, which better predict metabolic behavior, compared to traditional models that only consider stoichiometry and mass balance. However, adding and adjusting such constraints can represent a lengthy manual process. EMMAi relies on AI-predicted kinetic parameters to incorporate such constrains without manual intervention. Our results show that EMMAi ecGEMs lead to a significant increase in Pearson Correlation Coefficient (PCC) between predicted and experimentally measured growth rates in *Pseudomonas aeruginosa* PAO1 and *Escherichia coli* str. K-12 substr. MG1655. This improvement is especially pronounced in draft GEMs, where the introduction of enzyme constraints mitigates inaccuracies that potentially stem from incomplete curation and reaction gap-filling. By fully automating the integration of enzymatic constraints, EMMAi can improve the predictive accuracy of GEMs without the traditional burden of extensive manual refinement.

Traditional GEMs often require extensive manual curation to achieve reliable growth prediction accuracy. However, even curated models may not always provide optimal prediction accuracy. For example, recent studies have suggested that adding manually curated reactions can hinder model performance (13). Meanwhile, our results suggest that fully automated, accurate enzyme constraints serve as an effective layer of biological regulation that enhances model growth rate predictions without requiring manual curation. The increase in PCC for manually curated GEMs (average 0.27 improvement) and the drastic improvement in draft models (from −0.3 to 0.6) highlight the advantages of EMMAi’s approach. In doing so, EMMAi addresses the need for time-intensive manual adjustments. Unlike alternative ecGEM pipelines such as GECKO, AutoPACMEN, and ECMpy, which rely on experimentally measured or deep learning-predicted *kcat* values with limitations in prediction accuracy (21), EMMAi utilizes UniKP, an AI framework optimized for predicting kinetic parameters with a 13% improvement in correlation over DLKcat (22). This allows EMMAi to predict enzymatic constraints for a broader range of reactions, including those associated with non-model organisms.

One of the key methodological considerations in ecGEMs is the adjustment of the protein pool size. In our study, we adjusted protein pool constraints to match experimentally measured growth rates under a reference condition, as previously done in other enzyme-constrained models (14, 15). While the absolute values of protein pool sizes for the presented examples are outside the expected ranges, the fact that EMMAi achieves significant predictive improvements without requiring recalibration of *kcat* values is an important distinction that suggests a robust implementation of enzymatic constraints. Further, the improved performance of EMMAi-derived ecGEMs compared to a non-calibrated AutoPACMEN-derived (PCC increase of 0.22) suggests that high-quality *kcat* predictions translate directly into more accurate growth rate predictions.

Our findings underscore the potential of EMMAi as a powerful tool for expanding the applicability of GEMs in various research fields. The increased predictive accuracy of carbon-source-dependent growth rates positions EMMAi as a valuable asset for metabolic engineering applications, where precise prediction of growth rates is crucial (31). GEMs have been applied to study the roles of less-characterised organisms as part of microbiomes (4, 32, 33). The ability to generate ecGEMs without extensive manual intervention will improve the reliability of the predictions generated in such endeavours. Moreover, microbial consortia studies conventionally involve complex nutritional environments, where growth efficiency on available carbon source likely influence community interactions (34–36). Improved recapitulation of growth rates in GEMs will lead to the simulation of more realistic consortia dynamics. Therefore, the application of EMMAi carries the potential to accelerate research in microbiome studies, biotechnology, and synthetic biology.

Future work should explore benchmarking EMMAi across a broader set of organisms and metabolic conditions to validate its generalizability and identify potential areas for improvement. The application of AI-based kinetic parameter prediction to other aspects of constraint-based modeling, such as regulatory interactions, also presents an exciting avenue for future research. The field of AI is quickly evolving and models with improved prediction accuracy are likely to emerge soon. EMMAi’s structure allows for the coupling of alternatives to UniKP and future iterations of our software should test the reliability of such alternatives to produce improved GEMs based predictions.

Taken together, these results demonstrate that EMMAi represents a significant advancement in the field of metabolic modeling by providing a scalable and fully automated method for generating quality enzyme-constrained GEMs. By leveraging AI-driven kinetic predictions, EMMAi mitigates the need for labor-intensive manual curation while achieving superior predictive accuracy. As GEMs continue to be integral to systems biology and metabolic engineering, tools like EMMAi will play a crucial role in streamlining model development and enhancing their applicability to a wide range of biological questions.

## Supplementary material

### Supplementary file 1

Name: Supplementary file 1

Format: Excel (.xlsx)

Description: ecGEMs stoichiometric data, gene-protein-reaction associations and protein details for individual models in this study and smiles, gene sequences and predicted *kcat* value for individual reactions in models.

### Supplementary file 2

Name: Supplementary file 2

Format: Excel (.xlsx)

Description: Measured versus predicted growth rates for individual carbon sources across *E. coli* and

*P. aeruginosa* models discussed in this manuscript.

### Supplementary file 3

Name: Supplementary file 3

Format: Compressed directory (.zip)

Description: base GEMs, ecGEMs and implementation scripts.

## Acknowledgements

This work was supported by the Environment Business Unit; Data61 Business Unit; the Microbiomes for One Systems Health (MOSH)-Future Science Platform; the Advanced Engineering Biology (AEB)-Future Science Platform and a Scientific Computing Collaboration grant from the Commonwealth Scientific and Industrial Research Organisation.

