## Supplementary file 3 for "EMMAi: fast enzyme-allocation constraints in GEMs for improved biomass prediction across carbon sources": 1. data_retrieval.ipynb

```
# %% [code]
import os
import re
import cobra
import requests
import warnings
import pandas as pd
import pubchempy as pcp
from chemspipy import ChemSpider
from Bio import SeqIO
from Bio.SeqUtils import molecular_weight

warnings.filterwarnings(action='ignore', category=UserWarning)

strain = 'Pseudomonas aeruginosa (ATCC 15692, PAO1, ATCC 15692 / PAO1 / 1C / PRS 101 / LMG 12228)'
species = 'Pseudomonas aeruginosa'
model = cobra.io.read_sbml_model('test_PAO1/carved_GEM/carved_PAO1.xml')
model

# %% [code]
cs = ChemSpider('Iuwz5qtODtMS1s2Baf1Svu6sST0MzGdT')

def get_smiles_from_name(name):
    results = cs.search(name)
    if results:
        return results[0].smiles
    return "Compound not found"

def remove_characters_within_brackets(text):
    # Pattern to match content within brackets (including the brackets themselves)
    pattern = r'\s*\([^)]*\)'
    # Replace matched content with an empty string
    cleaned_text = re.sub(pattern, '', text)
    return cleaned_text

# %% [code]
def get_sequence_and_mass(protein_faa_path, gene_name):
    for record in SeqIO.parse(protein_faa_path, "fasta"):
        if gene_name in record.description:
            sequence = str(record.seq)
            mass = molecular_weight(sequence, seq_type='protein')
            return sequence, mass
    return None, None

# %% [code]
def get_accession(sp, gene):
    query = f"{gene} {sp}"
    uniprot_query = f'https://rest.uniprot.org/uniprotkb/stream?fields=accession%2Cec%2Cmass%2Cgene_names%2Clineage%2Corganism_name%2Csequence&format=tsv&query=%28{query}%29'
    uniprot_data = requests.get(uniprot_query).text.split('\n')
    data_df = pd.DataFrame([x.split('\t') for x in uniprot_data])
    data_df = data_df.rename(columns=data_df.iloc[0]).drop(data_df.index[0])
    for index, row in data_df.iterrows():
        if f'{sp}' in str(row['Organism']) and gene in str(row['Gene Names']):
            accession = row['Entry']
            mass = row['Mass']
            ec = row['EC number']
            seq = row['Sequence']
            return accession, mass, ec, seq
    return None, None, None, None

# extra = ['umpH', 'ldtB', 'ldtD', 'ldtC', 'pfo', 'glsB',
#          'ldtE', 'ldtA', 'wbbH', 'wzxB', 'umpG', 'fau',
#          'gpmM'
#         ]

# %% [code]
"""Gene-sequence retrieval from UniProt"""

# Paths
gene_sequence_path = 'test_PAO1/gene_sequence_data.csv'

# Initialize or load existing data
if os.path.exists(gene_sequence_path):
    genes_df = pd.read_csv(gene_sequence_path)
    processed_genes = set(genes_df['Gene name'].unique())
else:
    genes_df = pd.DataFrame(columns=['Gene ID', 'Gene name', 'Accession', 'Sequence', 
                                     'Mass', 'EC number', 'Organism', 'Gene reactions'])
    processed_genes = set()

# Initialize a list to collect updates
batch_updates = []
batch_size = 15

for gene in model.genes:
    if gene.name not in processed_genes:
        try:
            organism = strain
            accession, mass, ec, seq = get_accession(strain, gene.name)
            if seq == None:
                organism = species
                accession, mass, ec, seq = get_accession(species, gene.name)
            batch_updates.append(
                {'Gene ID': gene.id, 'Gene name': gene.name, 'Accession': accession,
                 'Sequence': seq, 'Mass': mass, 'EC number': ec, 'Organism': organism,
                 'Gene reactions': [r.id for r in gene.reactions]})
            processed_genes.add(gene.name)

            # Check if batch size is reached
            if len(batch_updates) >= batch_size:
                # Convert batch updates to DataFrame
                batch_df = pd.DataFrame(batch_updates)
                # Append batch updates to the main DataFrame and save
                genes_df = pd.concat([genes_df, batch_df], ignore_index=True)
                genes_df.to_csv(gene_sequence_path, index=False)
                # Clear batch updates after saving
                batch_updates = []

        except Exception as e:
            print(f"Error processing {m.id}: {e}")
            # Handle error (e.g., log it, attempt to recover, skip this metabolite, etc.)

# After the loop, check if there are any remaining updates not yet saved
if batch_updates:
    batch_df = pd.DataFrame(batch_updates)
    genes_df = pd.concat([genes_df, batch_df], ignore_index=True)
    genes_df.to_csv(gene_sequence_path, index=False)

print(f"Updated genes data saved to '{gene_sequence_path}'.")

# %% [code]
"""Gene-sequence retrieval from fasta file"""
protein_faa_path = 'test_PAO1/carved_GEM/protein.faa'

# Paths
gene_sequence_path = 'test_PAO1/gene_sequence_data.csv'

# Initialize or load existing data
if os.path.exists(gene_sequence_path):
    genes_df = pd.read_csv(gene_sequence_path)
    processed_genes = set(genes_df['Gene name'].unique())
else:
    genes_df = pd.DataFrame(columns=['Gene ID', 'Gene name', 'Accession', 'Sequence', 'Mass', 'EC number', 'Organism', 'Gene reactions'])
    processed_genes = set()

# Initialize a list to collect updates
batch_updates = []
batch_size = 15

for gene in model.genes:
    gene_id = gene.id.replace('_', '.')
    if gene_id not in processed_genes:
        try:
            # Protein.faa file reading and molecular weight calculation
            seq, mass = get_sequence_and_mass(protein_faa_path, gene_id)
            if seq is not None:
                batch_updates.append(
                    {'Gene ID': gene_id, 'Gene name': gene_id, 'Accession': None,
                     'Sequence': seq, 'Mass': mass, 'EC number': None, 'Organism': None,
                     'Gene reactions': [r.id for r in gene.reactions]})  # Update 'Gene reactions' as needed
                processed_genes.add(gene_id)

                # Check if batch size is reached
                if len(batch_updates) >= batch_size:
                    # Convert batch updates to DataFrame and append to main DataFrame
                    batch_df = pd.DataFrame(batch_updates)
                    genes_df = pd.concat([genes_df, batch_df], ignore_index=True)
                    genes_df.to_csv(gene_sequence_path, index=False)
                    batch_updates = []

        except Exception as e:
            print(f"Error processing {gene_id}: {e}")
            # Handle error

# Check for any remaining updates not yet saved
if batch_updates:
    batch_df = pd.DataFrame(batch_updates)
    genes_df = pd.concat([genes_df, batch_df], ignore_index=True)
    genes_df.to_csv(gene_sequence_path, index=False)

print(f"Updated genes data saved to '{gene_sequence_path}'.")

# %% [code]
"""Metabolite-SMILES retrieval"""

# Paths
metabolite_smiles_path = 'test_PAO1/metabolite_smiles_data.csv'

# Initialize or load existing data
if os.path.exists(metabolite_smiles_path):
    metabolites_df = pd.read_csv(metabolite_smiles_path)
    processed_metabolites = set(metabolites_df['metabolite_id'].unique())
else:
    metabolites_df = pd.DataFrame(columns=['metabolite_id', 'name', 'smiles'])
    processed_metabolites = set()

# Initialize a list to collect updates
batch_updates = []
batch_size = 15
API_counter = 0
API_test = '10-Formyltetrahydrofolate'

for m in model.metabolites:
    if m.id not in processed_metabolites:
        name = remove_characters_within_brackets(m.name)
        if m.formula is not None and m.formula in name:
            name = name.replace(m.formula, '')
        try:
            # Looking for corresponding metabolite smiles in ChemSpider and PubChem
            compounds = pcp.get_compounds(name, 'name')
            if len(compounds) > 0:
                for com in compounds:
                    smiles = com.isomeric_smiles
                    break
            else:
                smiles = get_smiles_from_name(name)
                API_counter += 1
                if API_counter == 1000:
                    print('SpiderChem API key no longer valid')
                    print(f'Test: {get_smiles_from_name(API_test)}')
                    
            # Collect data for batch update
            batch_updates.append({'metabolite_id': m.id, 'name': m.name, 'smiles': smiles})
            processed_metabolites.add(m.id)

            # Check if batch size is reached
            if len(batch_updates) >= batch_size:
                # Convert batch updates to DataFrame
                batch_df = pd.DataFrame(batch_updates)
                # Append batch updates to the main DataFrame and save
                metabolites_df = pd.concat([metabolites_df, batch_df], ignore_index=True)
                metabolites_df.to_csv(metabolite_smiles_path, index=False)
                # Clear batch updates after saving
                batch_updates = []

        except Exception as e:
            print(f"Error processing {m.id}: {e}")
            # Handle error (e.g., log it, attempt to recover, skip this metabolite, etc.)

# After the loop, check if there are any remaining updates not yet saved
if batch_updates:
    batch_df = pd.DataFrame(batch_updates)
    metabolites_df = pd.concat([metabolites_df, batch_df], ignore_index=True)
    metabolites_df.to_csv(metabolite_smiles_path, index=False)

print(f"Updated metabolites data saved to '{metabolite_smiles_path}'.")


# %% [code]
"""
Pair sequences with adequate substrate SMILES
"""

# metabolites that should not be considered main substrates of reactions
cofactors = [
    'ATP C10H12N5O13P3', 'H+', 'Nicotinamide adenine dinucleotide phosphate - reduced', 'H2O H2O', 'Appolipoprotein', 'CMP C9H12N3O8P',
    'Sodium', 'Nicotinamide adenine dinucleotide', 'Nicotinamide adenine dinucleotide - reduced', 'O2 O2', 'FMN C17H19N4O9P', 'Ubiquinol-8',
    'Coenzyme A', 'Acetyl-CoA','GTP C10H12N5O14P3', 'Flavin adenine dinucleotide oxidized', 'Flavin adenine dinucleotide reduced', 'UDP C9H11N2O12P2',
    'Acyl carrier protein', 'AMP C10H12N5O7P', 'ITP C10H11N4O14P3', "Pyridoxal 5'-phosphate", 'Nicotinamide adenine dinucleotide phosphate',
    'Reduced FMN', 'Reduced riboflavin', 'Iron (Fe3+)', 'Fe2+ mitochondria', 'Phosphate', 'Ubiquinone-8', 'Menaquinone 8', 'Ammonium', 'CO2 CO2', 
    'IMP C10H11N4O8P', 'Bicarbonate', 'Reduced glutathione', 'Tungsten binding cofactor', 'UTP C9H11N2O15P3', 'Hydrogen sulfide', 
    'Menaquinol 8', 'Reduced thioredoxin', 'Hydrogen peroxide', 'Nitric oxide', 'Hydrogen', 'ADP C10H12N5O10P2', 'Diphosphate', 'UMP C9H11N2O9P', 
    'GMP C10H12N5O8P', 'Riboflavin C17H20N4O6', 'GDP C10H12N5O11P2'
]

metabolite_smiles_path = 'test_PAO1/metabolite_smiles_data.csv'
metabolites_df = pd.read_csv(metabolite_smiles_path, index_col='metabolite_id')
gene_sequence_path = 'test_PAO1/gene_sequence_data.csv'
genes_df = pd.read_csv(gene_sequence_path, index_col='Gene ID')

seqs_smiles = []

for gene in model.genes:
    if gene.id == 'spontaneous':
        continue
    gene_id = gene.id.replace('_', '.')
    sequence = genes_df.loc[gene_id, 'Sequence']
    mass = genes_df.loc[gene_id, 'Mass']
    ec = genes_df.loc[gene_id, 'EC number']
    for r in gene.reactions:
        reactants = [(m.name, m.id) for m in r.reactants if m.name not in cofactors]
        for i in reactants:
            smiles = metabolites_df.loc[i[1], 'smiles']
            seqs_smiles.append({
                'Gene ID': gene_id, 'Gene name': gene.name, 'Sequence': sequence, 
                'Reaction ID': r.id, 'Reaction name': r.name, 'Reaction': r.reaction, 'Direction': 'Forward', 
                'Substrate Name': i[0], 'Substrate ID': i[1], 'Substrate Smiles': smiles, 'Kcat': ''
            })
        
        if r.reversibility:
            products = [(m.name, m.id) for m in r.products if m.name not in cofactors]
            for i in products:
                smiles = metabolites_df.loc[i[1], 'smiles']
                seqs_smiles.append({
                    'Gene ID': gene_id, 'Gene name': gene.name, 'Sequence': sequence, 
                    'Reaction ID': r.id, 'Reaction name': r.name, 'Reaction': r.reaction, 'Direction': 'Reverse', 
                    'Substrate Name': i[0], 'Substrate ID': i[1], 'Substrate Smiles': smiles, 'Kcat': ''
                })
seqs_smiles_df = pd.DataFrame(seqs_smiles)
seqs_smiles_df.to_csv('test_PAO1/sequences_smiles.csv', index=False)
```
