## Supplementary file 3 for "EMMAi: fast enzyme-allocation constraints in GEMs for improved biomass prediction across carbon sources": 2. uni_kp_prot.ipynb

```
# %% [code]
import torch
import warnings
from build_vocab import WordVocab
from pretrain_trfm import TrfmSeq2seq
from utils import split
# build_vocab, pretrain_trfm, utils packages are from SMILES Transformer
from transformers import T5EncoderModel, T5Tokenizer
# transformers package is from ProtTrans
import re
import gc
import numpy as np
import pandas as pd
import pickle
import math

warnings.filterwarnings(action='ignore', category=UserWarning)

def smiles_to_vec(Smiles):
    pad_index = 0
    unk_index = 1
    eos_index = 2
    sos_index = 3
    mask_index = 4
    vocab = WordVocab.load_vocab('vocab.pkl')
    def get_inputs(sm):
        seq_len = 220
        sm = sm.split()
        if len(sm)>218:
            print('SMILES is too long ({:d})'.format(len(sm)))
            sm = sm[:109]+sm[-109:]
        ids = [vocab.stoi.get(token, unk_index) for token in sm]
        ids = [sos_index] + ids + [eos_index]
        seg = [1]*len(ids)
        padding = [pad_index]*(seq_len - len(ids))
        ids.extend(padding), seg.extend(padding)
        return ids, seg
    def get_array(smiles):
        x_id, x_seg = [], []
        for sm in smiles:
            a,b = get_inputs(sm)
            x_id.append(a)
            x_seg.append(b)
        return torch.tensor(x_id), torch.tensor(x_seg)
    trfm = TrfmSeq2seq(len(vocab), 256, len(vocab), 4)
    trfm.load_state_dict(torch.load('trfm_12_23000.pkl', map_location=torch.device('cpu')))
    trfm.eval()
    x_split = [split(sm) for sm in Smiles]
    xid, xseg = get_array(x_split)
    X = trfm.encode(torch.t(xid))
    return X


def Seq_to_vec(Sequence):
    for i in range(len(Sequence)):
        if len(Sequence[i]) > 1000:
            Sequence[i] = Sequence[i][:500] + Sequence[i][-500:]
    sequences_Example = []
    for i in range(len(Sequence)):
        zj = ''
        for j in range(len(Sequence[i]) - 1):
            zj += Sequence[i][j] + ' '
        zj += Sequence[i][-1]
        sequences_Example.append(zj)
    ###### you should place downloaded model into this directory.
    tokenizer = T5Tokenizer.from_pretrained("prot_t5_xl_uniref50", do_lower_case=False)
    model = T5EncoderModel.from_pretrained("prot_t5_xl_uniref50")
    gc.collect()
    print(torch.cuda.is_available())
    # 'cuda:0' if torch.cuda.is_available() else
    device = torch.device('cuda:0' if torch.cuda.is_available() else 'cpu')
    model = model.to(device)
    model = model.eval()
    features = []
    for i in range(len(sequences_Example)):
        print('For sequence ', str(i+1))
        sequences_Example_i = sequences_Example[i]
        sequences_Example_i = [re.sub(r"[UZOB]", "X", sequences_Example_i)]
        ids = tokenizer.batch_encode_plus(sequences_Example_i, add_special_tokens=True, padding=True)
        input_ids = torch.tensor(ids['input_ids']).to(device)
        attention_mask = torch.tensor(ids['attention_mask']).to(device)
        with torch.no_grad():
            embedding = model(input_ids=input_ids, attention_mask=attention_mask)
        embedding = embedding.last_hidden_state.cpu().numpy()
        for seq_num in range(len(embedding)):
            seq_len = (attention_mask[seq_num] == 1).sum()
            seq_emd = embedding[seq_num][:seq_len - 1]
            features.append(seq_emd)
    features_normalize = np.zeros([len(features), len(features[0][0])], dtype=float)
    for i in range(len(features)):
        for k in range(len(features[0][0])):
            for j in range(len(features[i])):
                features_normalize[i][k] += features[i][j][k]
            features_normalize[i][k] /= len(features[i])
    return features_normalize


# %% [code]
seqs_smiles_df = pd.read_csv('')

with open('UniKP for kcat.pkl', "rb") as f:
    model = pickle.load(f)

batch = 0
batch_len = 20

predicted_pair = {}

# Function to check if a pair of Substrate Smiles and Sequence has been assessed
def is_assessed(substrate_smiles, sequence):
    return (substrate_smiles, sequence) in predicted_pair

# Function to check if reaction is a transport reaction
transporters = ['transport', 'symporter', 'diffusion', 'antiport', 'tranposrt']
def contains_keywords(cell):
    return any(keyword.lower() in str(cell).lower() for keyword in transporters)

for index, row in seqs_smiles_df.iterrows():
    if type(row['Sequence']) != float and not contains_keywords(row['Reaction name']) and np.isnan(row['Kcat']):
        if row['Substrate Smiles'] != 'Compound not found':
            pair_key = (row['Substrate Smiles'], row['Sequence'])
            if is_assessed(*pair_key):
                seqs_smiles_df.at[index,'Kcat'] = predicted_pair[pair_key]
            
            else:
                print(row['Reaction ID'])
                seq_vec = Seq_to_vec([row['Sequence']])
                smiles_vec = smiles_to_vec([row['Substrate Smiles']])
                fused_vector = np.concatenate((smiles_vec, seq_vec), axis=1)
                pre_kcat = model.predict(fused_vector)
                kcat = [math.pow(10, pre_kcat[i]) for i in range(len(pre_kcat))]
                seqs_smiles_df.at[index, 'Kcat'] = kcat
                
                # Store the predicted kcat in the dictionary
                predicted_pair[pair_key] = kcat[0]
    batch += 1
    if batch == batch_len:
        seqs_smiles_df.to_csv('', index=False)
        batch = 0
seqs_smiles_df.to_csv('', index=False)

# %% [code]
import sklearn
print(sklearn.__version__)
```
