## Supplementary file 3 for "EMMAi: fast enzyme-allocation constraints in GEMs for improved biomass prediction across carbon sources": 3. model_modification_carved_PAO1.ipynb

```
# %% [code]
import cobra
import pandas as pd

GREEN = '\033[92m'
YELLOW = '\033[93m'

model = cobra.io.read_sbml_model('test_PAO1/carved_GEM/gapfillled_PAO1.xml')

# sol = model.optimize()
# model.summary(sol)

# %% [code]
transporters = ['transport', 'symporter', 'diffusion', 'antiport', 'tranposrt']
ecmodel = model.copy()

GREEN = '\033[92m'
YELLOW = '\033[93m'

# Function to check for keywords
def contains_keywords(cell):
    return any(keyword.lower() in str(cell).lower() for keyword in transporters)

reversible_count = 0

"""Addressing reaction reversibility"""
for reaction in ecmodel.reactions:
    if (reaction.reversibility and not contains_keywords(reaction.name) and reaction not in ecmodel.boundary):
        reversible_count += 1

        bwd = reaction.copy()
        bwd.id = reaction.id + '_rev'
        bwd.bounds = (-1000.0, 0.0)

        reaction.bounds = (0.0, 1000.0)
        reaction.id = reaction.id + '_fwr'

        ecmodel.add_reactions([bwd])

expected_total = len(model.reactions) + reversible_count
print(f'{GREEN}There are {len(model.reactions)} reactions in the original model')
print(f'{GREEN}of which {reversible_count} are reversible.')
print(f'{GREEN}Therefore, the expected number of reactions in the ecModel should be {expected_total}')

print(f'{YELLOW}The total number of reactions in the ecModel are {len(ecmodel.reactions)}')

# %% [code]
"""Breaking reactions into isozymes"""

ecmodel2 = ecmodel.copy()
ecmodel2.name = 'ecPAO1'

def split_reaction_by_gpr(base_model, reaction):
    # Extract the GPR rule of the reaction
    gpr_rule = reaction.gene_reaction_rule
    gpr_name = reaction.gene_name_reaction_rule
    # print(gpr_rule, gpr_name)
    genes = {g.id: g.name for g in reaction.genes}
    # print(genes)
    # Split the rule by 'or', considering the precedence of 'and'
    parts = [part.strip() for part in gpr_rule.split(' or ')]
    
    new_reactions = []
    
    for i, part in enumerate(parts):
        # Create a new reaction for each part
        new_reaction_id = f"{reaction.id}_iso{i+1}"
        new_reaction = cobra.Reaction(new_reaction_id)
        new_reaction.name = f"{reaction.name} iso{i+1}"
        new_reaction.add_metabolites(reaction.metabolites)
        new_reaction.subsystem = reaction.subsystem
        new_reaction.lower_bound = reaction.lower_bound
        new_reaction.upper_bound = reaction.upper_bound
        
        # Add new reaction to the model
        # Assign GPR rule considering 'and' connections within each part
        # print(part)
        gene_ids_in_part = [g_id for g_id in genes.keys() if g_id in part]
        new_reaction.gene_reaction_rule = ' and '.join(gene_ids_in_part)

        # Modify gene id and name for them to match original reactions in the model
        # as gene name is by default added as gene id when using reaction.gene_reaaction_rule
        base_model.add_reactions([new_reaction])
        # r = base_model.reactions.get_by_id(new_reaction_id)
        # print(r, [(g.id, g.name) for g in r.genes])
        new_reactions.append(new_reaction)

    # Remove the original reaction from the model
    ecmodel2.remove_reactions([reaction])
    
    return len(new_reactions)

isozymes = 0
for r in ecmodel.reactions:
    reaction = ecmodel2.reactions.get_by_id(r.id)
    if (not contains_keywords(reaction.name) and reaction not in ecmodel2.boundary 
        and 'or' in reaction.gene_name_reaction_rule):
        isozymes = isozymes + split_reaction_by_gpr(ecmodel2, reaction) - 1 
            
print(isozymes)
ecmodel2

# %% [code]
"""
MASS/KCAT PSEUDOMETABOLITES FOR RESOURCE USAGE
"""

import numpy as np
# ecmodel3 = ecmodel2.copy()

updated_sns = pd.read_csv('test_PAO1/sequences_smiles.csv', index_col='Gene ID')
gene_sequence_mass = pd.read_csv('test_PAO1/gene_sequence_data.csv', index_col='Gene ID')

with ecmodel2:
    usage = cobra.Metabolite(
        'usage',
        name='resource_usage_pseudometabolite',
        compartment='c'
    )
    
    ecmodel2.add_metabolites([usage])
    ecmodel2.add_boundary(ecmodel2.metabolites.get_by_id("usage"), type="demand")
    usage_reaction = ecmodel2.reactions.get_by_id('DM_usage')
    usage_reaction.bounds = (-0.1, 0.0)
    
    for reaction in ecmodel2.reactions:
        mass = 0.0
        direction = ''
        if reaction.lower_bound >= 0 and reaction.upper_bound > 0:
            direction = 'Forward'
        elif reaction.lower_bound < 0 and reaction.upper_bound <= 0:
            direction = 'Reverse'
        if (not contains_keywords(reaction.name) and reaction not in ecmodel2.boundary 
            and len([g for g in reaction.genes]) > 0):
            # print(reaction.id, reaction.reaction, [g.name for g in reaction.genes])
            if reaction.lower_bound < 0 and reaction.upper_bound <= 0:
                reactant_ids = [m.id for m in reaction.products]
            elif reaction.lower_bound >= 0 and reaction.upper_bound > 0:
                reactant_ids = [m.id for m in reaction.reactants]
            # print(reactant_ids)
            
            preliminary_kcats = {}
            for g in reaction.genes:
                g_id = g.id.replace('_', '.')
                if g_id in updated_sns.index:
                    subset = updated_sns.loc[g_id]
                    if isinstance(subset, pd.Series):
                        subset = pd.DataFrame([subset])
                    for index, row in subset.iterrows():
                        if (row['Substrate ID'] in reactant_ids and not np.isnan(row['Kcat']) and
                           row['Reaction ID'] in reaction.id and direction == row['Direction']):
                            mass += gene_sequence_mass.at[g_id, 'Mass'] 
                            if row['Substrate ID'] not in preliminary_kcats.keys():
                                preliminary_kcats.update({row['Substrate ID']: [row['Kcat']]})
                            else:
                                preliminary_kcats[row['Substrate ID']].append(row['Kcat'])
                else:
                    print(f'Gene {g.id} not found in seq-smiles relationship table.')
                        
            if len(preliminary_kcats) > 0:
                # print(preliminary_kcats)
                kcats = []
                for substrate in preliminary_kcats:
                    kcats.append(sum(preliminary_kcats[substrate])/len(preliminary_kcats[substrate])) 
                kcat = min(kcats) * 3600                     # convert kcat to a /h
                coefficient = (mass * 0.001)/kcat            # convert g/mol (Da) to g/mmol
                # print(kcat, mass, kcat/mass)
                
                if reaction.lower_bound < 0 and reaction.upper_bound <= 0:
                    reaction.add_metabolites({
                        usage: coefficient
                    })
                elif reaction.lower_bound >= 0 and reaction.upper_bound > 0:
                    reaction.add_metabolites({
                        usage: - coefficient
                    })
                # print(reaction)
            else:
                continue

    sol = ecmodel2.optimize()
    print(ecmodel2.summary(sol))
    cobra.io.write_sbml_model(ecmodel2, 'test_PAO1/output_GEMs/ec_PAO1_fixed.xml')
```
