## Supplementary file 3 for "EMMAi: fast enzyme-allocation constraints in GEMs for improved biomass prediction across carbon sources": 4. patching_models_carved_PAO1.ipynb

```
# %% [code]
import cobra
import pandas as pd
import os
import numpy as np

ec_model = cobra.io.read_sbml_model('test_PAO1/output_GEMs/ec_PAO1_fixed.xml')

# %% [code]
transporters = ['transport', 'symporter', 'diffusion', 'antiport', 'tranposrt']

cofactors = [
    'ATP', 'ADP', 'H+', 'Nicotinamide adenine dinucleotide phosphate - reduced', 'H2O', 'Applipoprotein', 'CMP C9H12N3O8P',
    'Sodium', 'Nicotinamide adenine dinucleotide', 'Nicotinamide adenine dinucleotide - reduced', 'O2', 'FMN', 'Ubiquinol-8',
    'Coenzyme A', 'Acetyl-CoA','GTP', 'Flavin adenine dinucleotide oxidized', 'Flavin adenine dinucleotide reduced', 'UDP C9H11N2O12P2',
    'Acyl carrier protein', 'AMP C10H12N5O7P', 'ITP C10H11N4O14P3', "Pyridoxal 5'-phosphate", 'Nicotinamide adenine dinucleotide phosphate',
    'Reduced FMN', 'Reduced riboflavin', 'Fe3+', 'Phosphate', 'Ubiquinone-8', 'Menaquinone 8', 'Ammonium', 'CO2', 'IMP C10H11N4O8P',
    'Bicarbonate', 'Reduced glutathione', 'Tungsten binding cofactor', 'UTP C9H11N2O15P3', 'Hydrogen sulfide', 'Menaquinol 8', 'Reduced thioredoxin',
    'Hydrogen peroxide', 'Nitric oxide', 'Hydrogen', 'ADP C10H12N5O10P2', 'Diphosphate', 'UMP C9H11N2O9P', 'GMP C10H12N5O8P', 'Riboflavin C17H20N4O6',
    'GDP'
]

acp_related = 0
cof_related = 0
missing = 0
included = 0

def contains_keywords(cell):
    return any(keyword.lower() in str(cell).lower() for keyword in transporters)

for reaction in ec_model.reactions:
    if (reaction not in ec_model.boundary and not contains_keywords(reaction.name)):
        if ('resource_usage_pseudometabolite' not in [m.name for m in reaction.metabolites]
            and len([g for g in reaction.genes]) > 0):
            if  'G_s0001' not in [g.name for g in reaction.genes]:
                # print(reaction.id, reaction.name, reaction.reaction, [g.name for g in reaction.genes])
                missing += 1
                if 'ACP_c' in reaction.reaction:
                    acp_related += 1

                reaction_cofactors = 0
                for m in reaction.metabolites:
                    if m.name in cofactors:
                        reaction_cofactors += 1

                if len([m for m in reaction.metabolites]) == reaction_cofactors:
                    print('Reaction made of cofactors only!')
                    cof_related +=  1

        if 'resource_usage_pseudometabolite' in [m.name for m in reaction.metabolites]:
            included += 1

missing, acp_related, cof_related, included

# %% [code]
"""Average reaction coefficient"""

usage_coefficients = []

for reaction in ec_model.reactions:
    if 'resource_usage_pseudometabolite' in [m.name for m in reaction.metabolites]:
        for m in reaction.metabolites:
            if m.id == 'usage':
                coef = reaction.metabolites[m]
                usage_coefficients.append(coef)

average_coef = sum(usage_coefficients)/len(usage_coefficients)
abs(average_coef)

# %% [code]
patched_model = ec_model.copy()

# %% [code]
usage = patched_model.metabolites.get_by_id('usage')

excluded_reactions = [
    'Biomass reaction',
    'R_ATPM'
]

with patched_model:
    for reaction in patched_model.reactions:
        if (reaction not in patched_model.boundary and not contains_keywords(reaction.name)
           and reaction.name not in excluded_reactions):
            if ('resource_usage_pseudometabolite' not in [m.name for m in reaction.metabolites]):
                # print(reaction.bounds)
                if reaction.lower_bound < 0 and reaction.upper_bound <= 0:
                    # print('reverse')
                    reaction.add_metabolites({
                        usage: abs(average_coef)
                    })
                if reaction.lower_bound >= 0 and reaction.upper_bound > 0:
                    # print('forward')
                    reaction.add_metabolites({
                        usage: - abs(average_coef)
                    })
                # print(reaction.name, reaction.reaction)

    cobra.io.write_sbml_model(patched_model, 'test_PAO1/output_GEMs/ec_gapfilled_PAO1.xml')
```
