## Supplementary file 3 for "EMMAi: fast enzyme-allocation constraints in GEMs for improved biomass prediction across carbon sources": 5. model_comparison_carved_PAO1.ipynb

```
# %% [code]
import cobra
import pandas as pd
import os
import numpy as np

base_model = cobra.io.read_sbml_model('test_PAO1/carved_GEM/gapfillled_PAO1.xml')
ec_model = cobra.io.read_sbml_model('test_PAO1/output_GEMs/ec_gapfilled_PAO1.xml')     # final ecGEM

# %% [code]
base_media = {
     'EX_cd2_e': 1000.0, 'EX_co2_e': 1000.0, 'EX_cobalt2_e': 1000.0, 'EX_cu2_e': 1000.0,
     # 'EX_fe2_e': 1000.0, 
    'EX_fe3_e': 1000.0, 'EX_h_e': 1000.0, 'EX_h2o_e': 1000.0,
     'EX_k_e': 1000.0, 'EX_mg2_e': 1000.0, 'EX_mn2_e': 1000.0, 
    # 'EX_n2_e': 1000.0,
     'EX_na1_e': 1000.0, 'EX_nh4_e': 1000.0, 'EX_o2_e': 1000.0, 'EX_pi_e': 1000.0,
     'EX_so4_e': 1000.0, 'EX_zn2_e': 1000.0, 'EX_ca2_e': 1000.0, 'EX_cl_e': 1000.0,
     'EX_glc__D_e': 1000.0
}

ec_model.medium = base_media

with ec_model:
    ec_model.medium = base_media
    ec_model.reactions.DM_usage.bounds = (-1.3, 0.0)
    sol = ec_model.optimize()
    print(ec_model.summary(sol))

# %% [code]
base_media = {
     'EX_cd2_e': 1000.0, 'EX_co2_e': 1000.0, 'EX_cobalt2_e': 1000.0, 'EX_cu2_e': 1000.0,
     # 'EX_fe2_e': 1000.0, 
     'EX_fe3_e': 1000.0, 'EX_h_e': 1000.0, 'EX_h2o_e': 1000.0,
     'EX_k_e': 1000.0, 'EX_mg2_e': 1000.0, 'EX_mn2_e': 1000.0, 
    # 'EX_n2_e': 1000.0,
     'EX_na1_e': 1000.0, 'EX_nh4_e': 1000.0, 'EX_o2_e': 1000.0, 'EX_pi_e': 1000.0,
     'EX_so4_e': 1000.0, 'EX_zn2_e': 1000.0, 'EX_ca2_e': 1000.0, 'EX_cl_e': 1000.0,
     # 'EX_glc__D_e': 10.0#, removed for testing of other carbon sources
}

sources = [
    'EX_4abut_e', 'EX_ac_e', 'EX_cit_e', 
    'EX_ala__D_e', 
    'EX_fru_e', 
    'EX_glc__D_e', 'EX_mnl_e', 'EX_man_e', 'EX_glcn_e', 'EX_glyc_e', 
    'EX_ala__L_e', 'EX_asp__L_e', 'EX_his__L_e', 'EX_lac__L_e', 'EX_leu__L_e', 
    'EX_orn_e', 'EX_phe__L_e', 'EX_pro__L_e', 'EX_ser__L_e', 'EX_akg_e', 
    'EX_ptrc_e', 
    'EX_succ_e', 
    'EX_tre_e', 'EX_asn__L_e', 
    # 'EX_maln_e', 
    'EX_cellb_e', 'EX_h2o_e', 'EX_glu__L_e', 'EX_thr__L_e', 
    # 'EX_btd__RR_e', 
    'EX_crn_e'
]

output_dict = {}
output_index = []

for source in sources:
    carbon_reaction = source
    r = base_model.reactions.get_by_id(carbon_reaction)
    carbon_source_name = r.name.replace(' exchange', '')
    output_dict.update({carbon_source_name: []})

def run_model(model, carbon_bound, id):
    with model:
        output_index.append(id)
        for source in sources:
            try:
                carbon_reaction = source
                r = model.reactions.get_by_id(carbon_reaction)
                media = base_media.copy()
                media.update({carbon_reaction: carbon_bound})
                model.medium = media
                # print(model.medium)
                sol = model.optimize()
                # print(r.name, sol.objective_value)
                carbon_source_name = r.name.replace(' exchange', '')
                output_dict[carbon_source_name].append(sol.objective_value)
                # print(cobra.flux_analysis.pfba(model))
            except:
                print(f'Not testing {source} as its exchange reaction was not found in model {id}.')


# ec_model.reactions.DM_usage.bounds = (-1800.0, 0.0)
# run_model(ec_model, 10, 'ec_model_10')

# ec_model.reactions.DM_usage.bounds = (-750.0, 0.0)
# run_model(ec_model, 1000, 'ec_model_1000')

# patched_ec_model.reactions.DM_usage.bounds = (-2400.0, 0.0)
# run_model(patched_ec_model, 10, 'patched_ec_model_10')

# patched_ec_model.reactions.DM_usage.bounds = (-1550.0, 0.0)
run_model(base_model, 10, 'base_model')

ec_model.reactions.DM_usage.bounds = (-1.3, 0.0)
run_model(ec_model, 1000, 'ec_model')
# run_model(patched_bounds_model, 10, 'patched_bounds_model')
# run_model(original_model, 10, 'original_model')

df = pd.DataFrame(output_dict, index=output_index)
df = df.transpose()
df
```
