## Supplementary file 3 for "EMMAi: fast enzyme-allocation constraints in GEMs for improved biomass prediction across carbon sources": 5. model_comparison_e_coli.ipynb

```
# %% [code]
import cobra
import pandas as pd
import os
import numpy as np

ec_model = cobra.io.read_sbml_model('e_coli/new_models/eciML1515.xml')                                                  # EC constrained with UniKP predictions
im = cobra.io.read_sbml_model('e_coli/pre_existing_models/iML1515_reference.xml')                                       # Base GEM (im stands for iML1515)
sm = cobra.io.read_sbml_model('e_coli/pre_existing_models/iJO1366_sMOMENT_2019_06_25_STANDARD_EXCHANGE_SCENARIO.xml')   # EC constrained with AUTOPACMEN pre-manual curation

# %% [code]
star_media = {
    'EX_ca2_e': 1000.0, 'EX_cbl1_e': 0.01, 'EX_cl_e': 1000.0, 'EX_co2_e': 1000.0, 'EX_cobalt2_e': 1000.0, 'EX_cu2_e': 1000.0, 
    'EX_fe2_e': 1000.0, 'EX_fe3_e': 1000.0, 'EX_h_e': 1000.0, 'EX_h2o_e': 1000.0, 'EX_k_e': 1000.0, 
    'EX_mg2_e': 1000.0, 'EX_mn2_e': 1000.0, 'EX_mobd_e': 1000.0, 'EX_nh4_e': 1000.0, 'EX_ni2_e': 1000.0, 'EX_o2_e': 1000.0, 
    'EX_pi_e': 1000.0, 'EX_so4_e': 1000.0, 'EX_zn2_e': 1000.0,
    'EX_glc__D_e': 1000.0
}

with ec_model:
    ec_model.medium = star_media
    ec_model.reactions.DM_usage.bounds = (-0.86, 0.0)
    sol = ec_model.optimize()
    print(ec_model.summary(sol))

# %% [code]
star_media = {
    'EX_ca2_e': 1000.0, 'EX_cbl1_e': 0.01, 'EX_cl_e': 1000.0, 'EX_co2_e': 1000.0, 'EX_cobalt2_e': 1000.0, 'EX_cu2_e': 1000.0, 
    'EX_fe2_e': 1000.0, 'EX_fe3_e': 1000.0, 'EX_h_e': 1000.0, 'EX_h2o_e': 1000.0, 'EX_k_e': 1000.0, 
    'EX_mg2_e': 1000.0, 'EX_mn2_e': 1000.0, 'EX_mobd_e': 1000.0, 'EX_nh4_e': 1000.0, 'EX_ni2_e': 1000.0, 'EX_o2_e': 1000.0, 
    'EX_pi_e': 1000.0, 'EX_so4_e': 1000.0, 'EX_zn2_e': 1000.0
    # 'EX_glc__D_e': 1000.0,    removed for testing of other carbon sources
}

sources = [
    'ac', 'glyc', 'akg', 'ala__L', 'pyr', 'fru', 'gsn', 
    'acgam', 'fum', 'rib__D', 'lac__L', 'glcn', 'sbt__D',
    'gam', 'mal__L', 'succ', 'malt', 'g6p', 'man', 'tre',
    'xyl__D', 'gal', 'glc__D', 'mnl'
]

output_dict = {}
output_index = []

for source in sources:
    carbon_reaction = f'EX_{source}_e'
    r = ec_model.reactions.get_by_id(carbon_reaction)
    carbon_source_name = r.name.replace(' exchange', '')
    output_dict.update({carbon_source_name: []})

def run_model(model, carbon_bound, id):
    with model:
        output_index.append(id)
        base_media = star_media
        for source in sources:
            try:
                carbon_reaction = f'EX_{source}_e'
                r = model.reactions.get_by_id(carbon_reaction)
                media = base_media.copy()
                media.update({carbon_reaction: carbon_bound})
                model.medium = media
                # print(model.medium)
                sol = model.optimize()
                # print(r.name, sol.objective_value)
                carbon_source_name = r.name.replace(' exchange', '')
                output_dict[carbon_source_name].append(sol.objective_value)
                # print(cobra.flux_analysis.pfba(model))
            except:
                print(f'Not testing {source} as its exchange reaction was not found in model {id}.')

ec_model.reactions.DM_usage.bounds = (-0.86, 0.0)
run_model(ec_model, 1000, 'ec_model')

run_model(im, 10, 'base_model')

run_model(star, 1000, 'eciJO_*')

df = pd.DataFrame(output_dict, index=output_index)
df = df.transpose()
df
```
